# Apples before the fall: does shape stability coincide with maturity?

**DOI:** 10.1101/2020.08.03.235010

**Authors:** Maria D Christodoulou, Alastair Culham

## Abstract

Fruit shape is the result of the interaction between genetic, epigenetic, environmental factors, and stochastic processes. As a core biological descriptor both for taxonomy and horticulture, the point at which shape stability is reached becomes paramount in apple cultivar identification, and authentication in commerce. Twelve apple cultivars were sampled at regular intervals from anthesis to harvest over two growing seasons. Linear and geometric morphometrics were analyzed to establish if and when shape stabilized and whether fruit asymmetry influenced this. Shape stability was detected in seven cultivars, four asymmetric and three symmetric. The remaining five did not stabilize. Shape stability, as defined here, is cultivar-dependent, and when it occurs, it is late in the growing season. Geometric morphometrics detected stability more readily than linear, especially in symmetric cultivars. Key shape features are important in apple marketing, giving the distinctness and apparent uniformity between cultivars expected at point of sale.

## Introduction

Shape development is a highly complex series of interacting molecular, biochemical and environmental pathways. At the starting point, shape development and diversity are controlled by genetics. For example, the tomato fruit shape, which has great diversity due to domestication and improvement processes, is controlled by four genes (Azzi et al., 2015). Two of these (*OVATE* and *SUN*) control fruit elongation whereas the other two (*FASCIATED* and *LOCULE NUMBER*) control locule numbers (Azzi et al., 2015). The combination of mutations in these four genes can result in the domesticated tomato shape ranging from spheroid to flat and obovoid (Monforte, Diaz, Cano-Delgado, & van der Knaap, 2014).

Aside from genetic effects, shape development is also impacted by phenotypic plasticity. Phenotypic plasticity is defined as the morphological, phenotypical or behavioral changes that are caused as a direct response to environmental stimuli (Price, Qvarnström, & Irwin, 2003). There are multiple examples of shape change as an environmental response in both plants and animals. In a study of *Datura wrightii* hort. ex Regel floral shape, phenotypic plasticity was demonstrated as a response to watering regime (Elle & Hare, 2002). In a common-garden experiment, plants that were watered twice a week had consistently longer corollas compared with plants that received only rainfall (Elle & Hare, 2002). In a range of ecotypes of *Arabidopsis thaliana* (L.) Heynh., leaf shape is reportedly altered when light levels are reduced, presenting smaller leaves and longer petioles (Tsukaya, 2006).

As shape is a major morphological descriptor of an organism, shape stability and by extension morphological stability, become fundamental biological concepts. This is even more so for cultivars which are distinguished by often very minor morphological differences. A key element of cultivar description is variation in size and shape. As opposed to species, cultivars have a very clear definition, a fact primarily due to the man-made nature of cultivars. Cultivars are described based on the selection of particular characters and these characters are required to be distinct, uniform and stable (Brickell et al., 2009). If a cultivar, therefore, is described based on particular morphological characters (which could include shape) then there is a requirement for stability for these characters. Differences in commercial value of cultivars within a species can be substantial which gives rise to the need for confident identification of individual cultivars (M. D. Christodoulou, Clark, & Culham, 2020; M.D. Christodoulou, Battey, & Culham, 2018).

Closer to the focus of this work, there have been a number of studies on the effect of environmental conditions on apple fruit. McKenzie (1971), in a general survey of apple fruit shape in New Zealand, recorded differences in shape between apples of the same cultivar grown in the north and south of New Zealand. In a study of light quality effect on ‘Golden Delicious’ apples, Noè and Eccher (1996) established that light levels and quality can affect both fruit shape and russeting. On the other hand, when Tromp (1990) studied the environmental effects on fruit shape of ‘Cox’s Orange Pippin’ using controlled environment rooms he found no significant differences. What makes this finding particularly interesting is that the environment was controlled for all the trees from anthesis onwards. It is therefore possible that the phenotypic plasticity recorded by McKenzie (1971) was due to environmental differences prior to anthesis. This is of interest because it has been recorded that the effect of the *OVATE* gene in tomatoes is established prior to anthesis (Monforte et al., 2014). Although it is not argued that the apple will have the exact same mechanisms as the tomato, both are climacteric domesticated fruit therefore similarities are possible. Since, the first whole genome iteration for apples was released (Velasco et al., 2010), characters such as ripening, pedigree origins, and leaf morphology have been studied, making the genetic basis of fruit shape an expected future topic of study by horticultural geneticists (Migicovsky, Li, Chitwood, & Myles, 2018; Muranty et al., 2020; Peace et al., 2019).

Even though phenotypic plasticity is well recorded there are also processes counteracting the environmental impact on the phenotype, specifically two buffering processes that promote phenotypic stability have been described (Willmore, Klingenberg, & Hallgrimsson, 2005). Canalization is described as the ability of an organism to buffer against both environmental and genetic variation in order to maintain phenotypic stability (Breuker, Patterson, & Klingenberg, 2006). On the other hand, developmental stability is the ability of an organism to buffer against random variation in order to maintain phenotypic stability (Willmore et al., 2005). The two mechanisms, although clearly defined, are difficult to isolate in development work. In a study of *Drosophila melanogaster* Meigen wing shape and size, Breuker et al. (2006) demonstrated the possibility that the two buffering pathways were fundamentally parts of a single process. Examples of developmental buffering have been described in both animals and plants. Tsukaya (2003) reported that in *A. thaliana*, manipulation of cell size and numbers in developing leaves was to a certain degree buffered in order to produce a stable phenotype. Manipulation of cell division rates in *Pelargonium* leaves demonstrated a mechanism where the faster dividing cells compensated for the slower dividing ones, resulting in a consistent shape (Day & Lawrence, 2000).

Traditionally, apple fruit shape was described by comparison with a standard geometric object (Bultitude, 1983). Examples of the possible shape categories included: oblong, conical or round (Clark & Cleal, 2005). This practice, which is still common in both identification keys and collection curation tools, aimed to summarize the extensive variety of shapes by grouping using a collection of predefined geometrical shapes (Sanders, 2010). In order to quantify shape, Westwood (1962) used the ratio between length and diameter (L/D) of the fruit. Using the L/D ratio as a proxy for shape, Westwood (1962) suggested that shape for apple cultivars stabilized between day 60 and day 100 from anthesis. The use of the L/D ratio has both benefits and limitations. By using measurements in two perpendicular axes, it facilitated the description of the overall shape in two dimensions. It also permitted the comparison between shapes, with ratios below 1 indicating a fruit that is wider than long and vice versa.

The description of ratios is easy to communicate, and intuitive but there are analytical concerns which arise from such use. First, a ratio of two normally distributed variables is not necessarily normally distributed (Atchley, Gaskins, & Anderson, 1976). This means that a ratio may not be suitable for analysis using parametric techniques, which assume normality. Second, ratio use is inherently paradoxical: if the variables in the ratio are unrelated, the ratio calculation creates a relationship (Curran-Everett, 2013). If, on the other hand, the two variables are related, the ratio calculation will only successfully demonstrate this relationship if it is linear and crosses the origin (D. A. Jackson, Harvey, & Somers, 1990). Third, ratio use can give rise to spurious correlations if the two variables used are both affected by a common confounding factor (Tu, Law, Ellison, & Gilthorpe, 2010). These three issues can be avoided, while still using length and diameter measurements as shape proxies, by performing an analysis of covariance (Tu et al., 2010).

In this work, we aim to investigate if and when fruit shape stabilizes for apple cultivars, whether the timings of this are cultivar-dependent, and if shape stability timings as described through linear morphometrics differ from those described through geometric morphometrics. These findings will have immediate commercial application to the marketing of apple cultivars harvested prior to full physiological maturity.

## Materials & Methods

### Sample collection

Twelve apple cultivars were sampled throughout the 2013 growing season at regular intervals, beginning at two weeks from flowering (anthesis) and ending when the fruit was deemed ripe for eating by the orchard’s pickers. Ten fruit per cultivar were collected at every sampling point, with the exception of the last harvest were 20 fruit were collected. Anthesis for the orchard in 2013 occurred on 10/06/2013. All fruit were collected from the National Fruit Collection, in Brogdale, Kent, UK. In 2014, six of the 12 cultivars were randomly selected to be resampled. Anthesis for 2014 occurred on 15/05/2014. A full list of cultivars and their sampling times are available in the Supplementary Materials.

### Morphometric Data Collection

Measurements of length and diameter (linear morphometrics), and geometric morphometrics for each sample were collected as described in Christodoulou et al. (2018). Geometric morphometrics were only collected for the 2013 season. The chosen landmarks and linear morphometrics are summarized in Figure 1.

**Figure 1:**
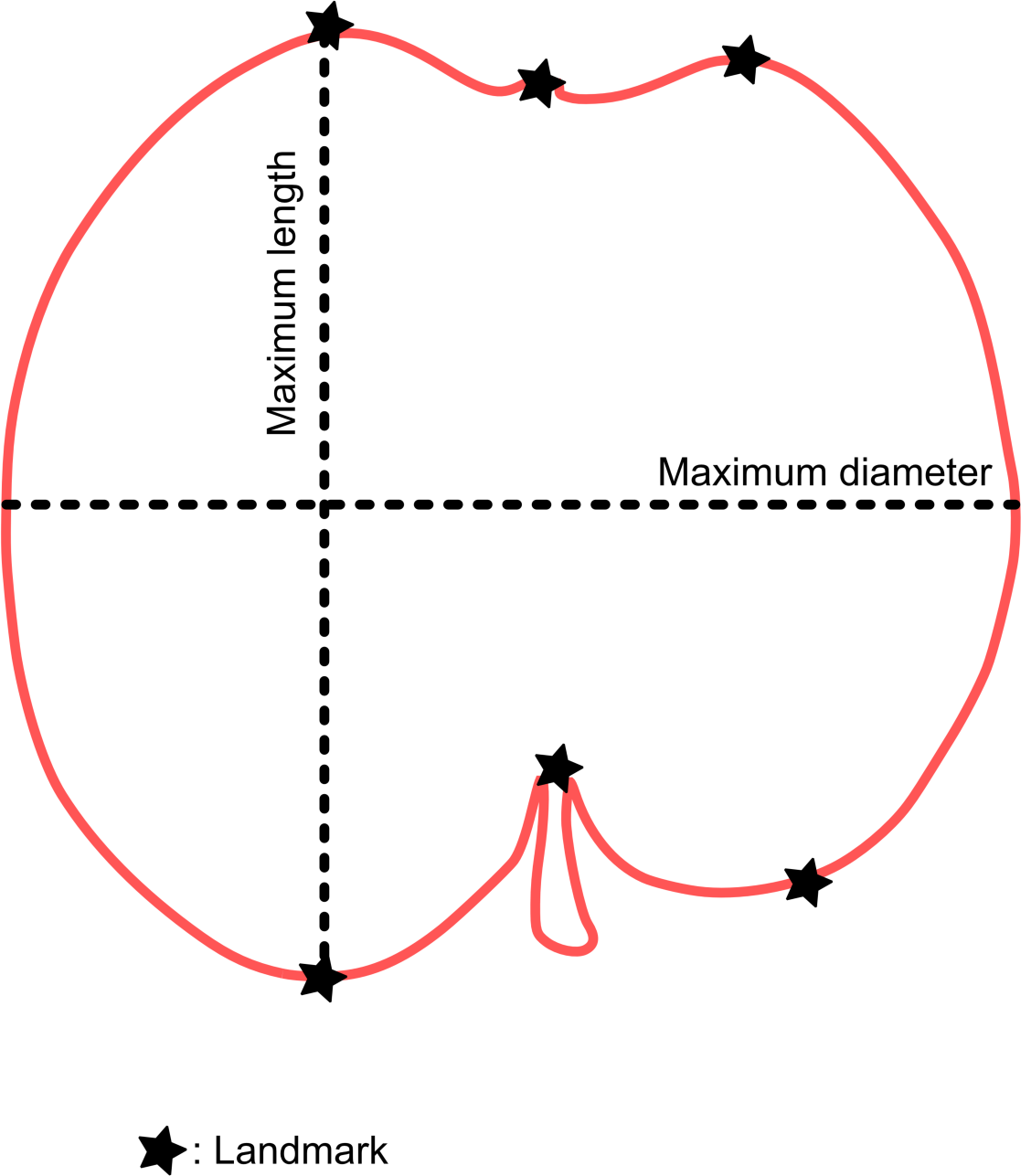
Digitised landmarks and linear morphometrics used on the sampled fruit. Six landmarks were selected per sample: two on the crown apices, one on the calyx, one on the pedicel attachment point, and two on the shoulder apices. Maximum length and diameter were measured using precision callipers.

### Shape Development Analysis

Analysis of Covariance (ANCOVA) was performed for each cultivar to examine how sampling week interacts with the correlation between length and diameter. To detect stability of shape for two sampling weeks, their slopes and intercepts had to be found not to be significantly different.

For the geometric morphometrics, regression for allometric effect was performed on MorphoJ (C. Klingenberg, 2011) and if found to be significant, analysis proceeded using the regressed dataset (C. P. Klingenberg & Marugán-Lobón, 2013; Openshaw & Keogh, 2014). If not, the dataset prior to allometric correction was employed. The allometric analysis step was performed to establish whether the effect of size was significant to the variation between samples. If the results were found to be significant, the dataset was corrected to exclude the variation that was explained by size. After removing the variation that was explained by size, the remaining variation between samples was attributed to shape differences. If the allometric analysis indicated that the effect of size was not significant for the variation between samples, the correction was not necessary and the data prior to regression were used. A Canonical Variates Analysis (CVA) was performed using harvest times as the classifier followed by a permutation test (10,000 permutations). The p-values of the Mahalanobis distances of the permutation test were used to establish shape differences between weeks (Zelditch, Swiderski, Sheets, & Fink, 2004).

### Asymmetry Analysis

To study whether differences in shape stability between cultivars were linked to fruit asymmetry, Procrustes ANOVAs were performed on the selected landmarks for the final harvest images for each cultivar (C. P. Klingenberg, Barluenga, & Meyer, 2002). In each case the axis of asymmetry was selected to pass through the calyx and pedicel landmarks. The two halves were compared with each other to establish whether there was statistical evidence of systematic asymmetry.

## Results

### Linear Morphometrics

Only three of the 12 cultivars stabilized prior to the last harvest. These were ‘Adam’s Pearmain’ (stable from Week 12 until Week 15), ‘Beacon’ (stable from Week 9 until Week 12), and ‘Wheeler’s Russet’ (stable from Week 12 until Week 17 in 2013, and Week 12 until 18 in 2014). All other cultivars demonstrated significant differences in the intercepts for the final sampling weeks. All code and model results are available as an R project, in the Supplementary materials.

### Geometric Morphometrics

When studied for allometry, 11 out of 12 cultivars demonstrated a significant allometric effect. The only cultivar that did not present a significant allometric effect was ‘Beacon’, and in this case the original data was not corrected for allometry. The remaining 11 cultivars were all corrected for allometry prior to further analysis. The percentage of variation explained by size as well as the associated p-values are summarized in Table 1.

**Table 1:** Cultivars that required allometric correction prior to analysis. All cultivars were regressed for allometric effect and 11 demonstrated significant effect of size (presented here) Each of the significantly affected cultivars is presented with variation explained and associated p-values.

| Cultivar | % of variation explained by size | p-value of 10,000 permutations test |
| --- | --- | --- |
| 'Adam's Pearmain' | 2.70% | 0.0003 |
| 'Boiken' | 3.92% | 0.0076 |
| 'Bovarde' | 4.07% | 0.0083 |
| 'Catshead' | 7.15% | <0.0001 |
| 'Fuji' | 3.44% | 0.0245 |
| 'Kaiser Franz Joseph' | 7.95% | <0.0001 |
| 'Limoncella' | 4.72% | 0.001 |
| 'Present van Engeland' | 23.25% | <0.0001 |
| 'Red Fortune' | 4.96% | 0.0109 |
| 'Rheinischer Krummstiel' | 6.00% | 0.0002 |
| 'Wheeler's Russet' | 3.05% | 0.0411 |

Each cultivar was studied using a CVA with harvesting times as the grouping factor. This was followed by a permutations test that compared the differences between all possible harvesting time combinations. The p-values from the 10,000 permutations test on the Mahalanobis distances were recorded for each harvest comparison. As the focus of this analysis was on the stability of shape between at least penultimate and ultimate harvests, only those comparison results are summarised in Table 2 below. Results from all other harvest comparisons can be found in the Supplementary Materials.

**Table 2:** Summary of Canonical Variates Analyses results between penultimate and ultimate harvests for the 12 studied cultivars. The p-values from the 10,000 permutations tests on the Mahalanobis distances are reported for each comparison (star significance in brackets).

| Cultivar | Penultimate Harvest Week | Final Harvest Week | p-value |
| --- | --- | --- | --- |
| 'Adam's Pearmain' | Week 12 | Week 15 | 0.0002 (***) |
| 'Beacon' | Week 9 | Week 12 | 0.0864 (NS) |
| 'Boiken' | Week 12 | Week 17 | 0.2119 (NS) |
| 'Bovarde' | Week 12 | Week 17 | 0.0069 (**) |
| 'Catshead' | Week 12 | Week 15 | 0.062 (NS) |
| 'Fuji' | Week 12 | Week 17 | 0.2973 (NS) |
| 'Kaiser Franz Joseph' | Week 12 | Week 15 | 0.0001 (***) |
| 'Limoncella' | Week 12 | Week 17 | 0.0057 (**) |
| 'Present van Engeland' | Week 12 | Week 15 | 0.0032 (**) |
| 'Red Fortune' | Week 9 | Week 12 | 0.0633 (NS) |
| 'Rheinischer Krummstiel' | Week 12 | Week 17 | 0.0093 (**) |
| 'Wheeler's Russet' | Week 12 | Week 17 | 0.1514 (NS) |

Shape stability for 2013, using both linear and geometric morphometrics, is summarised in Figure 2.

**Figure 2:**
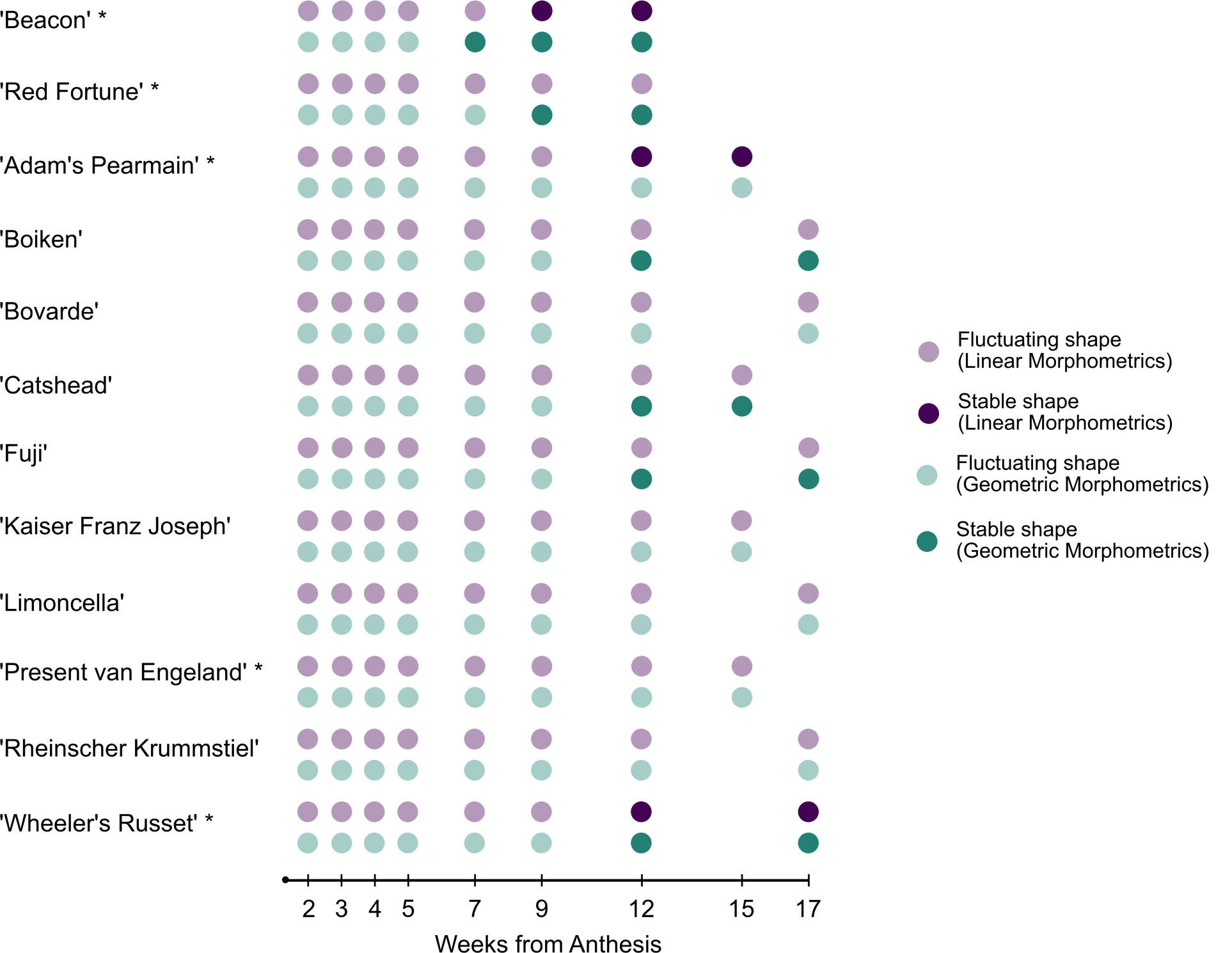
Shape stability status per Week/Cultivar for the 2013 growing season. Stability detection through linear morphometrics indicated in purple (pale for fluctuating shape, and dark purple for stable shape), and geometric morphometrics in green (pale for fluctuating shape, and dark green for stable shape). Asymmetric cultivars indicated with an asterisk after the cultivar name. Weeks are measured from flowering (anthesis). The two early season cultivars (‘Beacon’ and ‘Red Fortune’) are placed on the top of the chart. All other cultivars are main season cultivars.

Shape stability for the six cultivars repeated in 2014, using linear morphometrics, is summarised in Figure 3.

**Figure 3:**
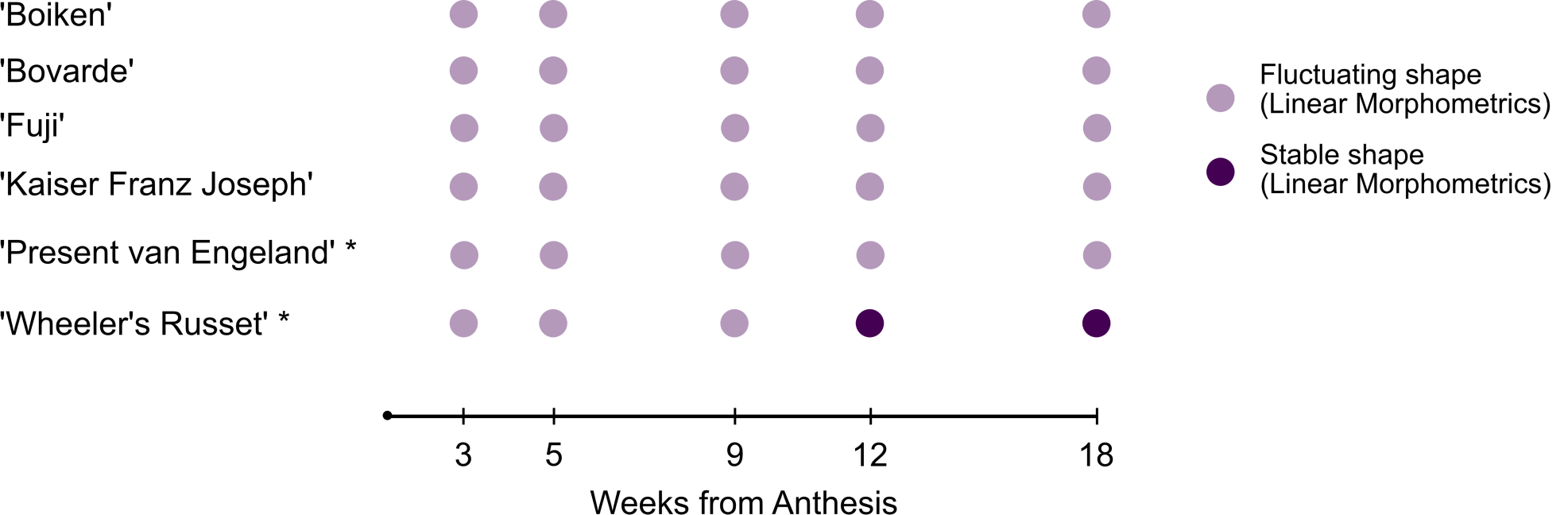
Shape stability status per Week/Cultivar for the 2014 growing season. Stability detection for that year was only conducted through linear morphometrics (pale purple for fluctuating shape, and dark purple for stable shape). Asymmetric cultivars indicated with an asterisk after the cultivar name. Weeks are measured from flowering (anthesis). All cultivars are main season cultivars.

The impact of the linear versus geometric morphometrics used to describe potential fruit shape stability can be observed in Figure 1. Whether stability happens and when it occurs matches on half of the tested cultivars for the two techniques. Five out of these six cultivars do not demonstrate shape stability as defined here (‘Bovarde’, ‘Kaiser Franz Joseph’, ‘Limoncella’, ‘Present van Engeland’, and ‘Rheinischer Krummstiel’). Shape stability using each method occurs at the same time only on ‘Wheeler’s Russet’. Cultivar ‘Beacon’ demonstrates shape stability for each of these methods but the point in time at which this is reached differs. For four of the remaining cultivars, stability is detected through geometric and not through linear morphometrics (‘Boiken’, ‘Catshead’, ‘Fuji, and ‘Red Fortune’). Shape stability was detected through linear morphometrics and not geometric only for ‘Adam’s Pearmain’.

Results from the symmetry analysis are in Table 3. Of the 12 cultivars, five were asymmetric and seven symmetric.

**Table 3:**
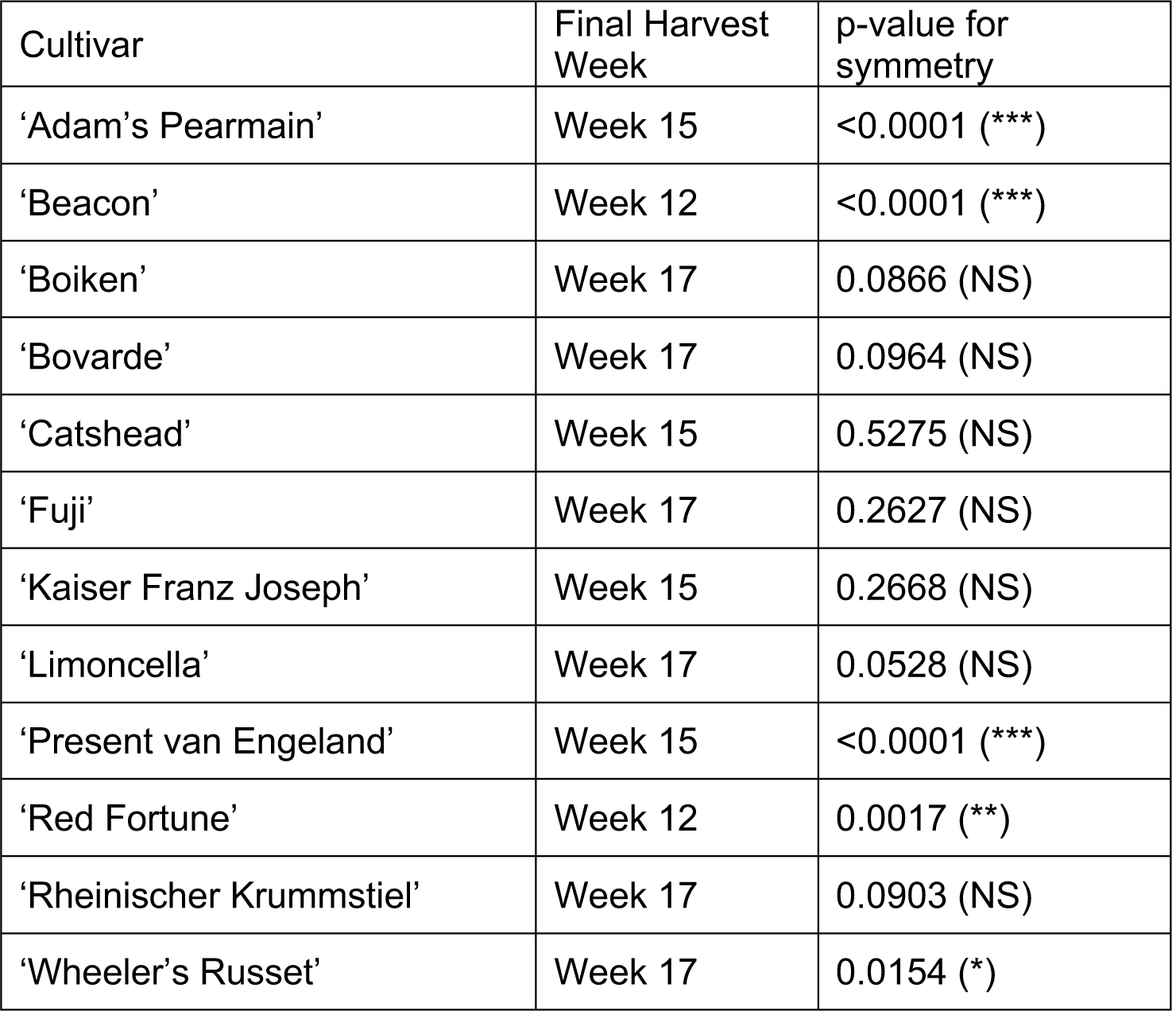
Symmetry analysis results for harvest week samples. Significant p-values (star significance in brackets), suggest substantial differences between the two sides of the fruit, indicating asymmetry.

Summary results for stability detection and presence of symmetry are in Table 4. Overall, presence of stability in symmetric cultivars was detected only through geometric morphometrics. For asymmetric fruit, both methods perform similarly.

**Table 4:** Summary table of stability detection and cultivar symmetry.

|  | Symmetric | Asymmetric |
| --- | --- | --- |
| No stability detection | 4 | 1 |
| LM first | 0 | 1 |
| GM first | 3 | 2 |
| Simultaneous detection | 0 | 1 |

## Discussion

Our results highlight two clear outcomes:

1. The method used to quantify shape impacts on our ability to describe and detect shape stability.
2. Shape stability, when it happens, occurs late in the season and is cultivar dependent.

The earliest time when stability is first detected in any cultivar is Week 7 (‘Beacon’, 49 days from anthesis), and the latest time of first detection is during Week 12 (e.g. ‘Catshead’, 84 days from anthesis). In total, seven of the 12 cultivars tested reach shape stability by at least one measure during this period. The remaining five cultivars do not achieve a stable shape by the time they are harvested based on our two approaches to measurement. This could be an accurate description of absence of stability or a consistent failure of either method to detect it. Comparing these findings with Westwood’s (1962) original observations of shape stability occurring between day 60 (Week 9) and day 100 (Week 14) from anthesis, suggests that for the majority of cultivars we are detecting similar trends. Whether these patterns are representative of shape stability is more a question of the reliability of our definition of the phenomenon than the presence of significant differences between weeks.

We define shape stability under linear morphometrics by testing whether the regression between length and diameter for the last harvest is significantly different to the regression of the preceding week. This therefore depends on whether we consider length and diameter in combination to be sufficient shape descriptors. For the geometric morphometrics, the method in combination with a permutation test, uses the position of the six landmarks selected to detect whether there are significant differences between weeks. Again, the success of the method relies on whether we believe the six landmarks selected to be sufficient shape descriptors. Both linear and geometric morphometrics, as described here, were used as underlying data for a machine learning classification method aiming to identify apple cultivars, which achieved 77.8% accuracy on an unclassified test set (Christodoulou, Battey & Culham, 2018). As such, we believe that the shape descriptions used for both methods are reflective of the cultivar shape, therefore their use for evaluation of shape stability is justified. Shape stability is an important commercial factor in fruit sales because customers expect uniform fruit at the point of purchase.

Linear morphometrics rely on individual measurements, geometric morphometrics combine positions of multiple landmarks. Comparison of findings between the two techniques illustrates this fundamental difference. In the case of shape stability this would be illustrated by geometric morphometrics detecting stability more readily than linear ones. This is the case for five cultivars in this study. Shape stability for symmetric apples was detected only using geometric morphometrics, whilst shape stability for asymmetric apples was detected in three out five cultivars by geometric morphometrics and two out of five by linear morphometrics. This means that linear morphometrics is not an effective approach to detect shape stability in symmetric apples and that geometric morphometrics is effective for both.

Some cultivars (such as ‘Beacon’ and ‘Wheeler’s Russet’) stabilised in shape prior to the final harvest week, while others did not appear to do so (e.g. ‘Bovarde’ and ‘Rheinischer Krummstiel’). This suggests that shape stability is cultivar dependent. These findings are congruent with the 2014 follow-up collection for limited samples. In terms of growing practices, the National Fruit Collection followed identical growing protocols between the two years. No differences in pest presence or management were recorded between the two years and pollination levels were similar. The observable difference in the two years is weather. Specifically, 2013 was cooler and drier than 2014. As this resulted in no differences in which cultivar stabilised prior to harvest, we can hypothesise that stability of shape is not easily perturbed by weather. We find that to be an unexpected finding, as it contradicted previous work. For example, McKenzie (1971) demonstrated that weather conditions can substantially affect shape in ‘Delicious’ apples grown in New Zealand. Studies on ‘Cox’s Orange Pippin’ suggested that cooler spring temperatures – such as the ones observed in 2013 - were associated with higher fruit yield by improving seed-set (J. E. Jackson & Hamer, 1980; J. E. Jackson, Hamer, & Wickenden, 1983). The fact that seed-set was affected by pre-anthesis temperatures is directly relevant to fruit shape as Drazeta et al. (2004) demonstrated increased asymmetry on ‘Granny Smith’ apples due to seed-set success and seed weight. These led us to expect differences in fruit shape stability between the two years which we did not observe. The levels of “June drop” (self-thinning immature fruit drop) were similar in the two years.

The observed shape stability differences between cultivars could be due to differences in developmental buffering. If the developmental stability or canalization abilities differed between cultivars, then some would be less prone to reaching a stable shape than others. It is therefore possible that the environmental conditions under which all the cultivars were grown, affected the developmental buffering mechanisms to different degrees, resulting in shape stability for some cultivars and none for others. Measuring developmental buffering in morphometric studies often relies on the concept of Fluctuating Asymmetry (FA) (C. P. Klingenberg et al., 2002). By measuring the differences between two structures in the same organism which are controlled by the same genetic mechanisms the stability of developmental process can be measured (Willmore et al., 2005). For example, if the asymmetry between the right wing and the left wing of a *Drosophila* is measured then the success of the developmental buffering for that *Drosophila* can be quantified. Since both wings were developed under exactly the same genetic, epigenetic and environmental conditions any differences between the two sides should be down to the buffering success of the organisms as a whole. Using FA as a measure of developmental buffering however may not be appropriate for apple fruit development. This is because the assumption of environmental stability between the two sides of the fruit may not hold. As apple fruit grow in clusters, the exposure of the individual fruit to the external environment varies. These different localised environmental conditions confound the developmental buffering. To measure developmental buffering for apples, we believe that a study needs to be conducted under strictly controlled environmental conditions, otherwise the environmental noise is likely to overcome the developmental signal.

We have previously shown that apples do not stop growing until they are harvested (Maria D Christodoulou & Culham, 2020). The mix of cultivars that show shape stability before harvest and those that don’t, may be indicative of growth and development of individual fruit slowing sufficiently resulting in our metrics not detecting subtle changes as the fruit approach physiological maturity. The evidence that changes become increasingly subtle as maturity is reached is congruent with the fact that apple cultivars can be recognised from their distinct shapes and sizes at harvest. Contrasting our findings with earlier work on authentication of apples using harvest shape (M.D. Christodoulou et al., 2018), we note that there is no correlation in the success in identification and the cultivars that stabilised prior to harvest.

## Conclusions

In this work, we aimed to establish if and when fruit shape of apple cultivars stabilizes prior to harvesting, and whether the findings are impacted by the method used - contrasting linear and geometric morphometrics. We established that the consistency of our findings is method-dependent, with geometric morphometrics detecting stability more readily than linear morphometrics. Some cultivars never reached stability prior to harvest. Whether or not the shape stabilized, apple cultivars are geometrically different from each other such that they can usually be identified by their shape. Remarkably, apple cultivars can be identified with a high degree of success despite continued growth and sometimes continued shape change up until the point of harvest.

## Acknowledgements

We thank Prof Nick Battey, Dr Matthew Ordidge, and the National Fruit Collection, Brogdale, Kent, for access to samples and assistance with sampling.

## Author contributions

MDC was responsible for data curation, formal analysis, methodology, validation, writing of manuscript. AC was responsible for conceptualization, funding acquisition, project administration, supervision, and writing of manuscript.

## Conflicts of Interest declarations in manuscripts

The authors declare no conflicts of interest.

## Data and Coding Availability

All data and code have been submitted as supplementary materials.

